# A Reproducible, Leakage-Aware Machine Learning Pipeline for Antimicrobial Peptide Activity Prediction

**DOI:** 10.64898/2026.11.26.769488

**Authors:** Isreal Abiodun Oluwafemi, Joshua Ampofo Yentumi

**Affiliations:** Department of Biochemistry, Olabisi Onabanjo University; Department of Biochemistry and Biotechnology, Kwame Nkrumah University of Science and Technology

**Keywords:** antimicrobial peptides, antimicrobial resistance, machine learning, feature engineering, ESMFold, SHAP, data leakage, reproducibility

## Abstract

Antimicrobial peptides (AMPs) are key alternatives to conventional antibiotics, but published machine learning models may overestimate predictive performance because of positive-class sequence redundancy and data leakage. We developed a leakage-aware pipeline using 13,469 peptides from DRAMP v5.0 and UniProt, clustered at 40% sequence identity with CD-HIT to enforce cluster-level splitting. A Random Forest trained on sequence and physicochemical features achieved an AUC-ROC of 0.939 and a Matthews correlation coefficient (MCC) of 0.671 on a cluster-disjoint test set (mean AUC-ROC 0.923 ± 0.012 across 20 repeated cluster-disjoint splits, versus 0.945 ± 0.004 under random splitting). Adding ESMFold-derived structural features did not improve aggregate metrics; a diagnostic analysis that separated directly folded from propagated sequences, and examined a length confound, showed a small positive difference (+0.010 AUC, not tested for significance) on directly folded sequences and a small negative difference (−0.007) on sequences with propagated features. SHAP analysis identified net cationic charge as the primary driver of predictions, consistent with known membrane-disruption mechanisms. Secondary classifiers of activity spectrum achieved AUC-ROCs of 0.871 (Gram-positive) and 0.851 (Gram-negative). Among 22 clinical-stage peptides with no exact-sequence match in the training or test sets, the mean predicted probability was 0.73; a high score for the non-antibacterial amphipathic peptide enfuvirtide suggests that the features capture broad membrane-interaction traits rather than antibacterial specificity. The pipeline is implemented as an open-source, 29-rule Snakemake workflow.

**Author Summary:** Bacteria are becoming resistant to antibiotics faster than new drugs are being developed, and this resistance already contributes to millions of deaths worldwide each year. Antimicrobial peptides are short protein fragments that form part of the innate immune defenses of many organisms, and they are among the most promising alternatives. There are far too many candidate peptides to test each one in the laboratory, so computer models that predict which peptides are likely to work could help researchers decide what to test first.

A hidden problem can make these models look better than they are. Public peptide databases contain many near-identical peptides, created when researchers change one or two residues in a peptide to study how it works. If near-duplicates end up in both the data a model learns from and the data used to test it, the model can score well simply by recognizing peptides it has effectively already seen. We built a prediction pipeline that groups similar peptides together before splitting the data, so that no near-duplicate appears on both sides.

Under this stricter test, a model based on peptide composition and chemistry still performed well, only about 0.02 AUC-ROC below its score under a conventional random split, and it relied mainly on electrical charge, which is consistent with how these peptides attack bacterial membranes. Adding predicted 3D structure did not improve results overall. A closer analysis suggested that the difference was largely linked to peptide length, and a small, untested possible benefit appeared on peptides whose structures were individually predicted. The model also scored a non-antibacterial membrane-active peptide highly, so its scores indicate membrane-interaction potential and not proven antibacterial activity. The complete pipeline is freely available so that other researchers can verify, reuse or extend it.

## Introduction

Antimicrobial resistance has moved from a projected threat to a measured one. The Global Research on Antimicrobial Resistance project estimated 4.71 million deaths associated with bacterial antimicrobial resistance in 2021, of which 1.14 million were directly attributable to it [1], extending the earlier global quantification for 2019 [2]. Forecasts to 2050 place the figures at 1.91 million attributable and 8.22 million associated deaths per year [1]. The same analysis shows the burden shifting toward older adults: deaths fell by more than half among children under five between 1990 and 2021 while rising by over 80% among adults aged 70 and above. Resistance is now a standing item on the global health agenda rather than an emerging concern [3].

The antibiotic development pipeline has not kept pace, which sustains interest in mechanistically distinct alternatives. Antimicrobial peptides (AMPs) are the most extensively studied of these. They are short, gene-encoded effectors of innate immunity, found across bacteria, plants, invertebrates, and vertebrates, and they have been characterized as an evolutionarily ancient and broadly conserved defense strategy [4]. Thousands of natural and synthetic sequences have since been catalogued and classified by source organism, structural fold, and activity spectrum [5]. Their appeal as therapeutics rests on a broad activity range that spans Gram-positive and Gram-negative bacteria, fungi, and enveloped viruses, together with immunomodulatory effects that conventional small-molecule antibiotics do not provide [6].

The dominant mechanism is physical rather than enzymatic. Most characterized AMPs are cationic and amphipathic. Electrostatic attraction to the anionic phospholipids and lipopolysaccharide of the bacterial envelope concentrates the peptide at the membrane surface, after which the hydrophobic face inserts into the lipid bilayer and compromises its integrity [7]. This coupling of charge and amphipathicity, rather than affinity for a single protein target, is what makes composition and physicochemical descriptors informative features for activity prediction.

Membrane disruption does not confer immunity from resistance, and the field has moved away from claiming that it does. Bacterial countermeasures are documented and include proteolytic degradation, surface-charge modification that reduces initial electrostatic capture, and efflux [8]. What distinguishes AMPs is the shape of the resistance problem rather than its absence: adaptation typically requires remodeling of conserved envelope architecture rather than a point mutation in a single drug target, and the resulting determinants are less readily transferred between species than the plasmid-borne genes that drive resistance to many conventional antibiotics [7, 8]. The practical implication is a slower and less mobile resistance trajectory, not an exemption from one.

That advantage is only useful at scale if promising sequences can be identified without screening each one at the bench, and the number of candidate sequences in public repositories now far exceeds what wet-lab characterization can absorb. Computational prediction is the standard response to that gap.

Machine learning has become the practical route to AMP discovery because wet-lab screening cannot keep pace with the number of candidate sequences already sitting in public databases such as Collection of Anti-Microbial Peptides (CAMPR3) [9] and Data Repository of Antimicrobial Peptides (DRAMP) [10]. Two broad design paradigms have emerged. The first treats AMP prediction as a classification problem over engineered descriptors, amino acid composition, physicochemical properties, and structural propensities, fed into standard classifiers such as random forests and support vector machines. iAMP-2L [11] and iAMPpred [12] established this approach, and AmPEP [13] refined it by mining distribution patterns of amino acid properties along the sequence rather than simple compositional counts. The appeal of this paradigm is transparency: a random forest built on physicochemical descriptors admits direct inspection of which properties drive a prediction, which matters for a field trying to connect predictions back to mechanism.

The second paradigm replaces engineered features with representations learned directly from sequence. Deep-AmPEP30 [14] applied a deep-learning approach to short peptides specifically, a length regime where compositional features are sparse and less informative. More recent work layers attention and multi-view architectures on top of learned representations: Pang et al. [15] combined a transformer encoder with imbalanced multi-label learning to predict activity type alongside AMP status, and Ghulam et al. [16] fused multiple feature views through a capsule network. These architectures trade the interpretability of the first paradigm for representational flexibility, and none of the studies we cite in either paradigm reports cluster-disjoint train/test splitting, although some may have reduced redundancy within their benchmarks by sequence-identity filtering before splitting, a related but different control that we did not assess for each study.

A separate line of work uses prediction not to classify a fixed set of candidates but to search or generate new ones. Huang et al. [17] mined the full space of possible short peptide sequences with a machine-learning pipeline rather than scoring a pre-existing library, and Bhangu et al. [18] paired a predictive gradient-boosted model with a generative component to propose novel candidates directly. Both point toward where the field is heading, from screening known sequences to designing new ones, but a generative model is only as trustworthy as the discriminator that scores its output, which returns the evaluation problem to center stage rather than past it.

That evaluation problem is more serious than the range of published accuracies suggests. Sidorczuk et al. [19] trained 660 models spanning twelve architectures and eleven negative-sampling strategies and cross-evaluated every combination. Performance was consistently inflated whenever a model’s training and benchmark negative sets were built by the same or a similar sampling method, meaning a large share of published AMP machine learning (AMP-ML) benchmarks compare a model against a version of itself rather than against an independent test of generalization. The authors concluded that it is not currently possible to say which published AMP predictor is the most accurate and built a dedicated benchmarking server, AMPBenchmark, specifically to give the field a fair comparison standard. Negative-class construction is not a footnote to AMP-ML methodology; it is a first-order determinant of what a reported number means.

The evaluation problem documented by Sidorczuk et al. [19] for negative-class construction has a positive-class counterpart that has received far less attention: sequence redundancy. AMP databases are curated as biological repositories, not machine-learning evaluation sets, and they accumulate large numbers of near-identical peptides from structure-activity-relationship studies, where a researcher systematically substitutes one or two residues in an otherwise fixed backbone to map which positions matter for activity. That curation logic is entirely reasonable for cataloguing biology. It is a problem for splitting data into train and test sets. If two near-duplicates from the same SAR series land on opposite sides of a split, a model can score well by recognizing a peptide it has effectively already seen, not by learning anything that generalizes to a genuinely novel sequence. A benchmark built this way is not measuring what it claims to measure, in much the same way that a negative set drawn from the same distribution as the training negatives is not measuring generalization either. The two problems share a root cause: split construction that does not respect what makes a test example genuinely independent of training.

This methodology aligns with recent moves toward stricter evaluation standards. Notably, Pham et al. [20] released GenPept-Curated-2025, a benchmark designed to enforce homology-controlled partitioning in AMP prediction, on the premise that conventional biological databases do not. Developed independently of the pipeline presented here, it reflects growing recognition that homology-aware evaluation matters for rigorous model assessment.

While Sidorczuk et al. [19] established that negative-class construction is a first-order determinant of a reported number, Pham et al. [20] and this paper together argue that positive-class redundancy is the other half of the equation. Neither concern is addressed by more data or a better classifier. Both require a different approach to splitting.

This paper’s approach is direct. Each class is clustered independently with Cluster Database at High Identity with Tolerance (CD-HIT) at 40% sequence identity and split train and test sets at the cluster level using StratifiedGroupKFold, treating cluster membership as the grouping variable rather than individual sequence identity. This is designed to ensure that no sequences placed in the same 40%-identity cluster, including near-duplicates and structure-activity-relationship (SAR) siblings, appear on both sides of the split. Zero cluster overlap between train and test is verified by a computational assertion prior to model training. Because CD-HIT is a heuristic clustering tool, this control is only as complete as the clustering itself, and more distant homologs may still span the split. We also quantify how much redundancy matters in practice by comparing random and cluster-disjoint splits of the same data.

A cluster-disjoint data split fundamentally alters the empirical questions that can be rigorously addressed by the data, directly motivating the second contribution of this study. Once sequence-level shortcuts are reduced, a critical question arises: does structural information beyond basic sequence and physicochemical composition provide incremental predictive value? Three-dimensional conformation serves as a natural candidate for evaluation, given that membrane-disrupting activity heavily depends on the amphipathic secondary structures a peptide adopts rather than its raw primary composition. Furthermore, because structure-derived features are not shared across the cluster-disjoint partitions, they cannot act as a leakage shortcut.

Because folding every individual sequence using ESMFold was computationally prohibitive under the available hardware constraints, a cluster-representative folding strategy was deployed. Specifically, three-dimensional structures were generated exclusively for the CD-HIT cluster representatives, and these extracted structural features were subsequently propagated to the remaining members of each respective cluster. This approach is a compromise dictated by computational tractability rather than an ideal design, so we also test whether it preserves a genuine structural signal or mainly introduces propagation noise.

## Materials and Methods

An overview of the workflow is shown in Fig. 1.

**Fig. 1.**
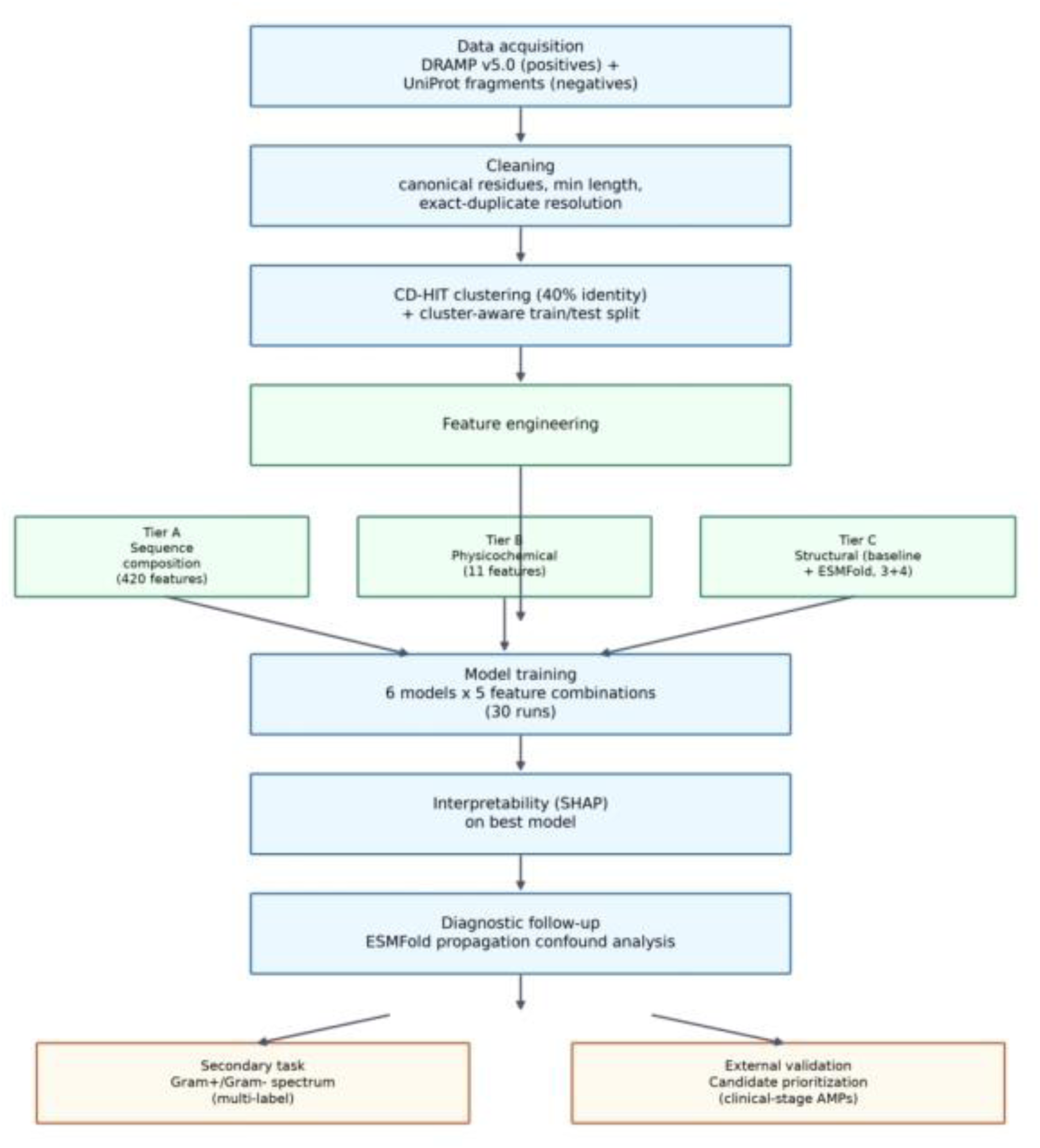
Pipeline overview: data acquisition through cleaning, clustering, feature engineering (three tiers), model training, interpretability, and two downstream validation tasks.

### Data Acquisition

Positive-class sequences were obtained from DRAMP v5.0 [10] (snapshot downloaded 2026-09-05; database lineage described in [21, 22]), yielding 12,784 raw entries in the general AMP category. DRAMP is updated regularly, and entries added after this date are not included. The snapshot’s SHA-256 checksum is recorded in the provenance manifest. Negative-class sequences were derived from the UniProtKB/Swiss-Prot database (accessed 2026-09-07; [23]): one random fragment was extracted from each of 3,383 reviewed, full-length proteins of at least 150 residues, with fragment length sampled from the cleaned positive-sequence length distribution under a fixed random seed (seed = 42) to prevent length itself from acting as a trivial class signal. Candidate proteins were excluded if their UniProt annotation matched any of a keyword list targeting antimicrobial-related function; antimicrobial, antibiotic, defensin, bacteriocin, antifungal, antiviral, antiparasitic, host defense, host-defense, membrane-active, toxic, cytotoxic, secretory, and lysozyme, to reduce the risk of drawing negatives from unannotated or cryptic antimicrobial sequences. One fragment containing a non-standard residue was subsequently removed, leaving 3,382 negative-class sequences.

This negative-sampling design falls into the random-fragment-extraction category evaluated by [19] rather than the whole-protein or motif-restricted alternatives also in use in the field. This approach was selected for two reasons specific to the goals of this pipeline, rather than as a claim that it resolves the sampling-bias problem documented by those authors in general. First, the fragment-length matching removes an obvious length confound between classes. Second, the annotation-keyword exclusion serves as a direct, auditable control against the most likely contamination route, namely reviewed proteins with real but undocumented antimicrobial activity.

This strategy does not eliminate the broader concern that any single negative-sampling method encodes assumptions about the operational definition of a non-AMP, a limitation addressed further in the Discussion section.

### Cleaning

Sequences were processed through a deterministic pipeline: uppercase normalization with a preserved flag for D-amino-acid residues (157 positive-class sequences flagged and retained through downstream steps, since D-stereochemistry does not itself invalidate standard-residue composition features), exclusion of sequences containing non-canonical residue codes, a minimum length threshold of 5 residues, and resolution of exact duplicates by preferring the entry with a valid database identifier, then alphabetically by identifier where no valid identifier existed. Non-canonical-residue exclusions were checked against DRAMP’s structured metadata rather than assumed to be curation artifacts; inspection confirmed the excluded sequences represent genuine synthetic modifications (including dimethylation, prenylation, and pyroglutamation, among others) that place them outside what a standard 20-amino-acid feature representation can capture.

Of the 12,784 raw positive-class sequences, 1,837 (14.4%) were excluded for non-canonical residues, 174 (1.4%) for falling below the 5-residue minimum, and 686 (5.4%) as exact duplicates, leaving 10,087 clean positive-class sequences (78.9% retention). Combined with the 3,382 negative-class sequences, the cleaned dataset used for all downstream modeling comprises 13,469 sequences. Full breakdown and cross-tabulation appear in Table 1.

**Table 1.** Data Cleaning Summary.

| **Step** | **Count** | **% of raw** |
|---|---|---|
| Raw input sequences | 12,784 | 100.0% |
| Excluded: non-canonical residues | 1,837 | 14.4% |
| Excluded: too short (<5 residues) | 174 | 1.4% |
| Excluded: exact duplicates | 686 | 5.4% |
| Retained | 10,087 | 78.9% |

*Note: 157 of the 10,087 retained sequences carry D-amino acid residues, flagged during normalization and preserved rather than excluded; this figure is not part of the exclusion accounting above.*

### Redundancy Control

Sequences were clustered independently within each class using CD-HIT [24] at a 40% sequence identity threshold (word size = 2). Crucially, the minimum sequence length parameter was relaxed (-l 4) to prevent CD-HIT’s default length cutoff from silently discarding short AMPs, several of which in the cleaned dataset fall within the 5–10 residue range. The 40% threshold was chosen deliberately as a permissive clustering criterion rather than a strict one: a high-identity threshold (90-100%), the conventional setting for exact-duplicate removal, only merges near-identical sequences and would leave structure-activity-relationship (SAR) variants (sequences differing at several residue positions but sharing a fixed backbone and near-identical function) free to land in different clusters and leak across a split. A 40% threshold pulls these more divergent but functionally redundant variants into a shared cluster, which should reduce that leakage path, although clustering at this threshold is not exhaustive for short peptides. This sits near the edge of CD-HIT’s recommended operating range; the tool’s documentation flags reduced reliability of its short-word filtering strategy below approximately 40% identity and recommends its slower sister tool, psi-cd-hit, for lower thresholds. Operating at the boundary of that range is a limitation of this clustering step, not a claim that redundancy control below 40% would be unnecessary.

Clustering was run independently on the positive and negative classes rather than jointly, as the two classes occupy distinct biological contexts. The positive class consists of curated antimicrobial sequences, on one side, while the negative class comprises randomly sampled fragments of the general reviewed proteome. Consequently, a shared clustering pass would impose a single similarity structure across sequence spaces with no meaningful shared ancestry. This yielded 885 positive-class clusters and 997 negative-class clusters.

MMseqs2 [25] was considered an alternative; it offers substantially faster clustering on very large sequence collections. CD-HIT was used instead because its explicit minimum-length control (-l) gives direct, transparent handling of the short-sequence tail characteristic of AMPs, which matters more for a 13,469-sequence dataset than raw clustering throughput does. This is a stated preference for this dataset’s scale and composition, not a general claim that CD-HIT outperforms MMseqs2.

### Splitting

Train/test partitioning used scikit-learn’s StratifiedGroupKFold [26] with cluster identity as the grouping variable and five splits, from which one group-disjoint fold was isolated as the held-out test set (representing a single cluster-disjoint train/test partition, not repeated cross-validation). This produced 10,775 training sequences (80.00%) and 2,694 test sequences (20.00%) (Table 2), with a hard assertion confirming zero cluster overlap between the two before any downstream step proceeded. Positive-class fractions were nearly identical between splits (0.7490 training, 0.7483 test) despite the whole-cluster constraint, indicating the class-stratification component of the split held despite the coarser grouping unit.

**Table 2.** Dataset and Split Summary.

|  | **Count** |
|---|---|
| Total sequences (positive + negative) | 13,469 |
| Positive clusters | 885 |
| Negative clusters | 997 |
| Training sequences | 10,775 |
| Test sequences | 2,694 |
| Cluster overlap between splits | 0 |

### Feature Engineering

Three feature tiers were computed for each sequence, corresponding to the combinations reported in Table 3.

**Table 3.** Top 10 Model/Feature Combination Results by AUC-ROC.

| Combination | Model | Features | AUC-ROC | AUPRC | F1 | MCC |
|---|---|---|---|---|---|---|
| seq+pcp | Random Forest | 431 | 0.9389 | 0.9799 | 0.9081 | 0.6713 |
| seq+pcp+struct(baseline) | LightGBM | 434 | 0.9382 | 0.9790 | 0.9158 | 0.6957 |
| seq+pcp+struct(baseline) | Random Forest | 434 | 0.9381 | 0.9797 | 0.9092 | 0.6742 |
| seq+pcp | LightGBM | 431 | 0.9369 | 0.9783 | 0.9175 | 0.6961 |
| seq+pcp | XGBoost | 431 | 0.9368 | 0.9785 | 0.9169 | 0.6887 |
| seq+pcp+struct(ESMFold) | Random Forest | 435 | 0.9348 | 0.9786 | 0.9058 | 0.6593 |
| all tiers | Random Forest | 438 | 0.9347 | 0.9786 | 0.9078 | 0.6643 |
| seq only | Random Forest | 420 | 0.9342 | 0.9779 | 0.9119 | 0.6856 |
| seq+pcp+struct(baseline) | XGBoost | 434 | 0.9305 | 0.9763 | 0.9089 | 0.6639 |
| seq only | XGBoost | 420 | 0.9274 | 0.9753 | 0.9086 | 0.6599 |

For the Sequence Composition Tier (Tier A), amino acid composition (20 features) and dipeptide composition (400 features) were computed directly from the primary sequence tokens, yielding 420 descriptors total.

While dipeptide composition captures local, first-order sequence adjacency information, this multi-tiered architecture deliberately excludes the long-range correlation vector mathematics characteristic of Pseudo-Amino Acid Composition (PseAAC) encodings [27], serving as a clean compositional baseline free of sequence-order-coupling parameter assumptions.

For the physicochemical tier (Tier B), eleven physicochemical descriptors, including net charge, charge density, isoelectric point, Boman index [28], and Eisenberg hydrophobic moment [29], were computed using the modlAMP package [30]. These descriptors assume a uniformly free C-terminus, an assumption that does not hold for all natural antimicrobial peptides (AMPs). Because C-terminal amidation is common among membrane-active peptides and shifts the terminal carboxyl group’s ionization state, charge and isoelectric-point values for amidated sequences are calculated as if the C-terminus were unblocked. This introduces a source of systematic, rather than random, error concentrated in a biologically identifiable subset of the dataset, representing a permanent metadata constraint imposed by the underlying source archive.

For the Structural Propensity Tier (Tier C), two variants were computed. The baseline variant used Biopython [31] to estimate helix, turn, and sheet propensity from sequence alone (3 features), providing a structural signal that requires no 3D coordinates. The ESMFold variant used predicted three-dimensional structures. For computational tractability under hardware constraints (a 2-core CPU with no GPU), full-atom folding via Meta’s ESM Atlas API [32] was run only on the 1,882 Cluster Database at High Identity with Tolerance (CD-HIT) cluster representatives, rather than on all 13,469 sequences individually. Backbone coordinates were extracted from the resulting structures with Biopython, and secondary structure was assigned from those coordinates using PyDSSP, a Python reimplementation of the DSSP algorithm [33].

This produced four structural features per representative sequence: fractional helix, sheet, and coil content, and mean per-residue pLDDT as a confidence measure. Each non-representative sequence within a cluster then inherited its representative’s four structural values unmodified. This direct propagation, rather than a re-estimation, ensures that a cluster’s non-representative members carry identical structural feature values to their representative and to each other. Five feature-tier combinations were evaluated in total: sequence alone (Tier A), sequence plus physicochemical (A+B), sequence plus physicochemical plus baseline structure (A+B+C-baseline), sequence plus physicochemical plus ESMFold structure (A+B+C-ESMFold), and all tiers combined (A+B+C-baseline+C-ESMFold).

### Model Training and Evaluation

Six classifiers were evaluated across the five feature-tier combinations described above (30 runs total): Logistic Regression, LinearSVC, Random Forest [34], XGBoost [35], LightGBM [36], and a multilayer perceptron (MLP). Logistic Regression, LinearSVC, Random Forest and the MLP were implemented in scikit-learn [26], while XGBoost and LightGBM were used through their own libraries. All models used a single fixed hyperparameter configuration across every feature-tier combination, chosen without any tuning search, to keep the comparison a clean ablation over feature tiers rather than confounding it with per-combination hyperparameter optimization. Random Forest, XGBoost, and LightGBM used 300 trees each, increased uniformly from each library’s default of 100 for ensemble stability rather than tuned per combination. Logistic Regression, LinearSVC, and the MLP required raising max_iter above default values to 2000, 5000, and 300, respectively, to ensure convergence; the MLP additionally used early stopping.

The scale-sensitive Logistic Regression, LinearSVC, and MLP models were trained on standardized features featuring zero mean and unit variance and fit on the training split only. Conversely, tree-based models utilized unstandardized features, consistent with their inherent scale invariance.

Class imbalance was addressed strictly within the training fold to avoid any test-set information influencing how the imbalance correction was calibrated. The balanced class weight parameter was applied to Logistic Regression, Random Forest, LightGBM, and the MLP, reweighting each class inversely proportional to its frequency in the training data. Because XGBoost does not support direct class weighting, it instead utilized the positive scale weight parameter set to the ratio of negative-to-positive sequence counts within the training split. This split contained 2,704 negative and 8,071 positive training sequences, yielding a positive scale weight value of approximately 0.335. Model evaluation utilized four metrics computed on the held-out, cluster-disjoint test set: the Area Under the Receiver Operating Characteristic (AUC-ROC) curve, the Area Under the Precision-Recall Curve (AUPRC), the F1-score, and the Matthews Correlation Coefficient (MCC). F1-score and MCC both require a hard classification decision and were computed at the default 0.5 probability threshold, with no post-hoc threshold tuning performed on the test fold. This protocol ensures the reported MCC (0.6713) and F1-score (0.9081) for the top model remain an unbiased reflection of default-threshold performance rather than an artifact optimized against the evaluation set. AUC-ROC was treated as the primary ranking criterion for identifying the optimal model architecture, given that it evaluates prediction ranking quality independent of a classification threshold; F1-score and MCC are reported as supplementary rather than primary selection metrics.

### Statistical Testing

Uncertainty on the best model’s test-set metrics was estimated by bootstrap resampling: the test set was resampled with replacement 1,000 times (fixed seed = 42), and AUC-ROC and MCC were recomputed on each resample to build empirical 95% confidence intervals (2.5th/97.5th percentiles). To test whether the top entries in Table 3 are distinguishable from one another, a paired bootstrap was used: the same resampled test-set indices were applied across all compared models within each of the 1,000 iterations, so each comparison uses matched data rather than independent resamples, and a 95% confidence interval was computed on the paired AUC-ROC difference for each pair. A difference was treated as statistically significant only if its interval excluded zero. These intervals reflect resampling of test sequences only; because test sequences within a cluster are not independent, they are conditional on this single partition and are likely narrower than the true uncertainty (see Sensitivity Analyses and Discussion). Classifier and feature-tier selection used test-set AUC-ROC, and no separate validation set was held out, so the reported performance of the selected model is subject to optimistic selection bias.

The bootstrap resampling script used for confidence interval and significance estimation was drafted with assistance from Claude (Anthropic). The script was executed exclusively by the authors on their own trained models and held-out test data; no data, predictions, or results were generated by the AI tool. All reported values were independently cross-checked by the authors against the pipeline’s saved outputs (model_results.csv, and related files) before inclusion in this manuscript.

### Sensitivity Analyses

To quantify how much sequence redundancy inflates performance, the Random Forest on sequence and physicochemical features was trained and evaluated on 20 repeated random splits (20% test, stratified by class, ignoring cluster membership) and 20 repeated cluster-disjoint splits (StratifiedGroupKFold, five folds, one fold held out), using seeds 42 to 61. Cluster identifiers were prefixed with the class label, and zero cluster overlap was asserted for every cluster-disjoint split. As baselines, raw net charge was used directly as a score without a model, and a logistic regression was trained on net charge, charge density and isoelectric point under the same cluster-disjoint splits.

### Interpretability

Feature attributions for the best-performing model (Random Forest on sequence composition and physicochemical features) were computed with SHAP (SHapley Additive exPlanations) TreeExplainer using interventional feature perturbation [37, 38]. The direction of each feature’s effect was defined as the Spearman rank correlation between the feature’s value and its SHAP value across the 200 explained sequences; a positive correlation means higher feature values push the prediction toward the antimicrobial class.

The background distribution for the explainer was a 100-sequence sample drawn from the training split, and attributions were computed on a 200-sequence subsample of the test set; both samples were drawn without replacement by deterministic random sampling under the project’s fixed seed (42), without class stratification. Fixing the seed and sampling procedure here, as elsewhere in the pipeline, means the reported SHAP values (Table 4) are exactly reproducible from the same code and data rather than dependent on an unlogged random draw. Because attributions were computed on only 200 test sequences, near-tied feature ranks should not be over-interpreted.

**Table 4.** Top 10 Features by Mean Absolute SHAP Value.

| **Rank** | **Feature** | **Mean \|SHAP\|** | **Direction (Spearman ρ)** |
|---|---|---|---|
| 1 | pcp_Charge | 0.0595 | + (0.91) |
| 2 | pcp_ChargeDensity | 0.0387 | + (0.95) |
| 3 | aac_C (cysteine) | 0.0319 | + (0.47) |
| 4 | pcp_pI (isoelectric point) | 0.0208 | + (0.92) |
| 5 | aac_E (glutamate) | 0.0203 | - (-0.85) |
| 6 | pcp_uH_eisenberg (hydrophobic moment) | 0.0184 | + (0.86) |
| 7 | pcp_BomanInd | 0.0150 | - (-0.59) |
| 8 | aac_K (lysine) | 0.0149 | + (0.87) |
| 9 | aac_D (aspartate) | 0.0120 | - (-0.80) |
| 10 | dpc_KK | 0.0118 | + (0.80) |

### Reproducibility

The complete pipeline, from raw data download through final figures, is implemented as a Snakemake workflow [39] comprising 29 rules organized into modular functional tracks: data acquisition and cryptographic checksum validation, sequence normalization and filtration, class-independent CD-HIT clustering, multi-tiered feature engineering (compositional, physicochemical, baseline structural, and ESMFold-derived structural propagation), parallel model training and ablation across the six classifiers and five feature-tier combinations, interpretability profile generation, and automated validation-set scoring. Each rule declares its inputs and outputs explicitly, so the full dependency graph and any downstream step affected by a change upstream is recoverable directly from the workflow definition rather than from prose description. A version-controlled data-provenance manifest records the source URL, download timestamp, and SHA-256 checksum for every raw data file, and the fixed seed of 42 is used consistently across negative-class fragment sampling, train/test splitting, and SHAP background/evaluation sampling.

The full pipeline was executed and benchmarked on a commodity Ubuntu Linux machine with a dual-core Intel Core i5 processor, 16 GB of RAM, and no GPU acceleration, the same hardware constraint that motivated the cluster-representative folding strategy in the structural feature tier (see “ESMFold Structural Features: A Diagnostic Case Study” in Results). The full software dependency stack, including exact package versions for scikit-learn, XGBoost, LightGBM, modlAMP, Biopython, and SHAP, is frozen in a version-controlled environment.yml file distributed with the pipeline, so the computational environment itself is reproducible, not only the code.

## Results

### Dataset Construction and Cleaning

DRAMP v5.0 provided 12,784 raw positive-class sequences [10]. Cleaning removed 1,837 sequences (14.4%) containing non-canonical residue codes, 174 (1.4%) below the 5-residue minimum length, and 686 (5.4%) as exact duplicates, leaving 10,087 clean positive-class sequences (78.9% retention; Table 1).

The non-canonical-residue exclusions were not treated as presumed artifacts. Verification against the structured metadata of DRAMP, rather than an assumption of sequencing noise, revealed that these sequences represent genuine synthetic modifications, including dimethylated, prenylated, and pyroglutamated residues. These modifications place the sequences outside what a standard 20-amino-acid feature representation can capture, making their exclusion a representational necessity rather than a data-quality correction. Separately, 157 positive-class sequences carrying D-amino acid residues were flagged during normalization and retained rather than excluded (Table 1). Because D-stereochemistry does not by itself compromise standard-residue composition features, these sequences remain in the modeling dataset with their stereochemical status preserved for any future feature tier.

The duplicate-resolution step surfaced a specific data-quality issue: one entry’s identifier had been corrupted into the literal string “nan” within the DRAMP export. This anomaly was detected because it caused a downstream join operation to fail, necessitating a targeted inspection of the affected row rather than allowing a silent join failure to discard the sequence without documentation.

This single case highlights the structural requirement that automated cleaning pipelines incorporate explicit validation checkpoints designed to surface anomalies transparently rather than executing silent operations.

Negative-class sequences were built by fragment extraction from reviewed UniProt proteins, yielding 3,382 negatives from 3,383 source proteins after excluding one fragment containing a non-standard residue (full extraction and annotation-filtering parameters are given in Methods).

Both classes were clustered independently with CD-HIT at 40% identity, producing 885 positive clusters and 997 negative clusters, and split at the cluster level with StratifiedGroupKFold, with a hard assertion confirming zero cluster overlap between train and test.

This produced 10,775 training sequences and 2,694 test sequences, with positive-class fractions nearly identical between splits (0.7490 training, 0.7483 test) despite the coarser, whole-cluster grouping constraint (Table 2).

As an early verification step prior to modeling, the mean net charge was compared between classes, yielding +2.73 for positives and -0.39 for the negative class. This distribution aligns with the expected direction for the conventional cationic-membrane mechanism, providing a critical confirmation that the physicochemical feature computation functioned correctly prior to downstream applications.

### Model Ablation: Sequence, Physicochemical, and Structural Features

Six classifiers, including Logistic Regression, LinearSVC, Random Forest, XGBoost, LightGBM, and a Multilayer Perceptron (MLP), were trained across five feature-tier combinations for a total of 30 runs (Fig. 2). The top 10 results by AUC-ROC are reported in Table 3.

**Fig. 2.**
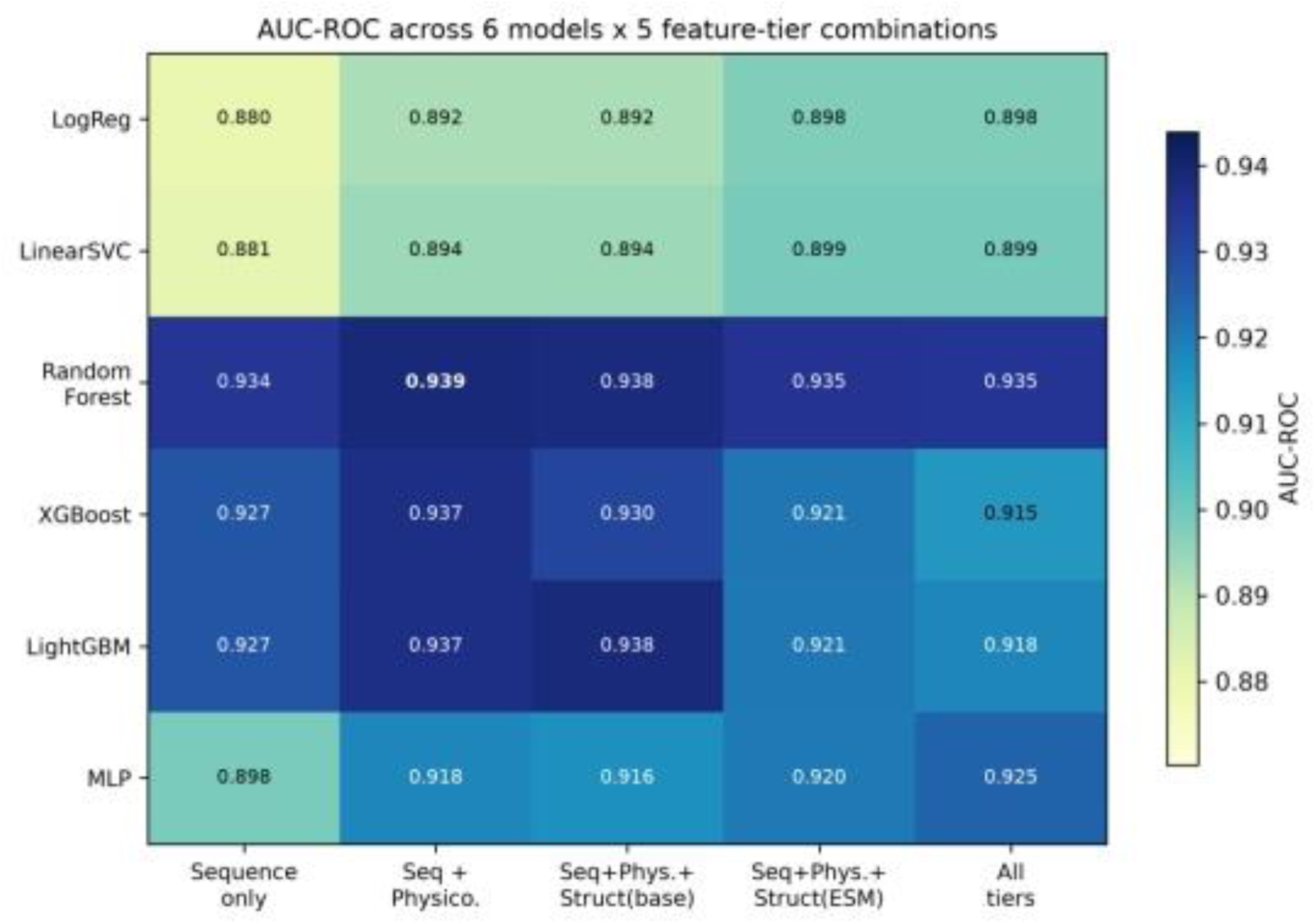
AUC-ROC heatmap across all 6 models x 5 feature-tier combinations

The best result came from Random Forest on sequence composition plus physicochemical features (431 features): AUC-ROC 0.9389 (95% CI: 0.9308-0.9477, bootstrap resample of the test set, 1,000 iterations), AUPRC 0.9799, F1 0.9081, MCC 0.6713 (95% CI: 0.6426-0.7071).

The top five results in Table 3 span a narrow band from 0.9389 to 0.9368 AUC-ROC, representing a spread of roughly two-tenths of a point, and all five utilize tree-based ensemble architectures, specifically Random Forest, LightGBM, and XGBoost. This performance plateau was tested directly with a paired bootstrap comparison among the top three combinations over 1,000 resamples of the test set. The same resampled indices were applied across all compared models within each iteration to ensure comparisons were matched rather than independent. None of the three pairwise AUC-ROC differences were statistically significant at the 95% confidence level. Specifically, the seq+pcp Random Forest versus seq+pcp+struct LightGBM yielded a mean difference of +0.0008 (95% CI: [-0.0048, +0.0060]), while the comparison against seq+pcp+struct Random Forest showed a difference of +0.0009 (95% CI: [-0.0012, +0.0028]). Similarly, seq+pcp+struct LightGBM versus seq+pcp+struct Random Forest resulted in a mean difference of +0.0001 (95% CI: [-0.0055, +0.0057]).

Metric selection within this performance band is non-neutral. On the sequence composition plus physicochemical feature set specifically, LightGBM yields a higher F1-score (0.9175 versus 0.9081) and MCC (0.6961 versus 0.6713) than Random Forest; these threshold-dependent differences were not tested for significance. That different metrics favor different models within a band of AUC-ROC values that the paired bootstrap could not separate is consistent with a performance plateau. These results suggest that, once sequence composition and physicochemical descriptors are captured, the tree-based models perform similarly on this cluster-disjoint split, and we found no evidence that classifier choice within the top tier matters. Absence of a significant difference is not proof of equivalence, and only the top three combinations were compared.

Physicochemical features provided a modest, untested improvement in AUC-ROC over sequence composition alone for Random Forest, raising the AUC-ROC from 0.9342 to 0.9389. Neither structural tier improved on the best sequence and physicochemical combination; the baseline structural tier changed AUC-ROC by −0.0008 for Random Forest and +0.0013 for LightGBM. Furthermore, the ESMFold tier specifically underperformed the sequence and physicochemical features across most classifiers; for example, XGBoost performance decreased from 0.9368 to 0.9211, and LightGBM dropped from 0.9369 to 0.9208, an unexpected degradation that warrants systematic diagnostic investigation.

### ESMFold Structural Features: A Diagnostic Case Study

An unexpected outcome of this magnitude required diagnostic investigation rather than simple reporting. The initial hypothesis was that because only 14% of the sequences possessed an individually computed fold (with the remaining 86% inheriting a cluster representative’s structural values via propagation due to computational hardware constraints), this propagation mechanism was introducing substantial noise into the models. To evaluate this hypothesis, the test set was partitioned into 376 sequences containing directly computed structural features and 2,318 sequences that inherited features via propagation.

The resulting performance metrics did not support the initial noise-dilution hypothesis. The directly folded representative subset achieved a lower average performance (0.863 AUC) than the propagated subset (0.918 AUC). If feature propagation were purely introducing random noise, the representative subset containing genuine structural information should have demonstrated superior predictive performance. This inverse relationship suggested that an unmodeled confounding factor contributed to the performance discrepancy.

Deeper analysis revealed that representative sequences are systematically longer (mean 47.5 residues) than propagated sequences (mean 22.4 residues). The default behavior of CD-HIT is to designate the longest sequence in each cluster as its representative, a mechanical side effect of the clustering algorithm that may carry an independent, non-structural relationship with classification difficulty. Consequently, a subset selected by the criterion of possessing an individual fold is, by virtue of how CD-HIT selects representatives, also a subset selected for greater sequence length, making these two properties inseparable by construction.

To isolate which factor was responsible for the performance difference, a control experiment was executed using the same representative versus propagated split but excluding all ESMFold features from the models. This control experiment revealed a gap of similar size (a decrease of 0.072 AUC, with representatives scoring below propagated sequences) compared to the ESMFold-inclusive baseline (a decrease of 0.055 AUC). Because the performance gap persisted when the structural features were removed entirely, it cannot be attributed to the ESMFold features or to their propagation. Sequence length, which differs systematically between the two subsets, is a plausible driver of the gap, but we did not test this directly, for example by length-stratified analysis (Fig. 3).

**Fig. 3.**
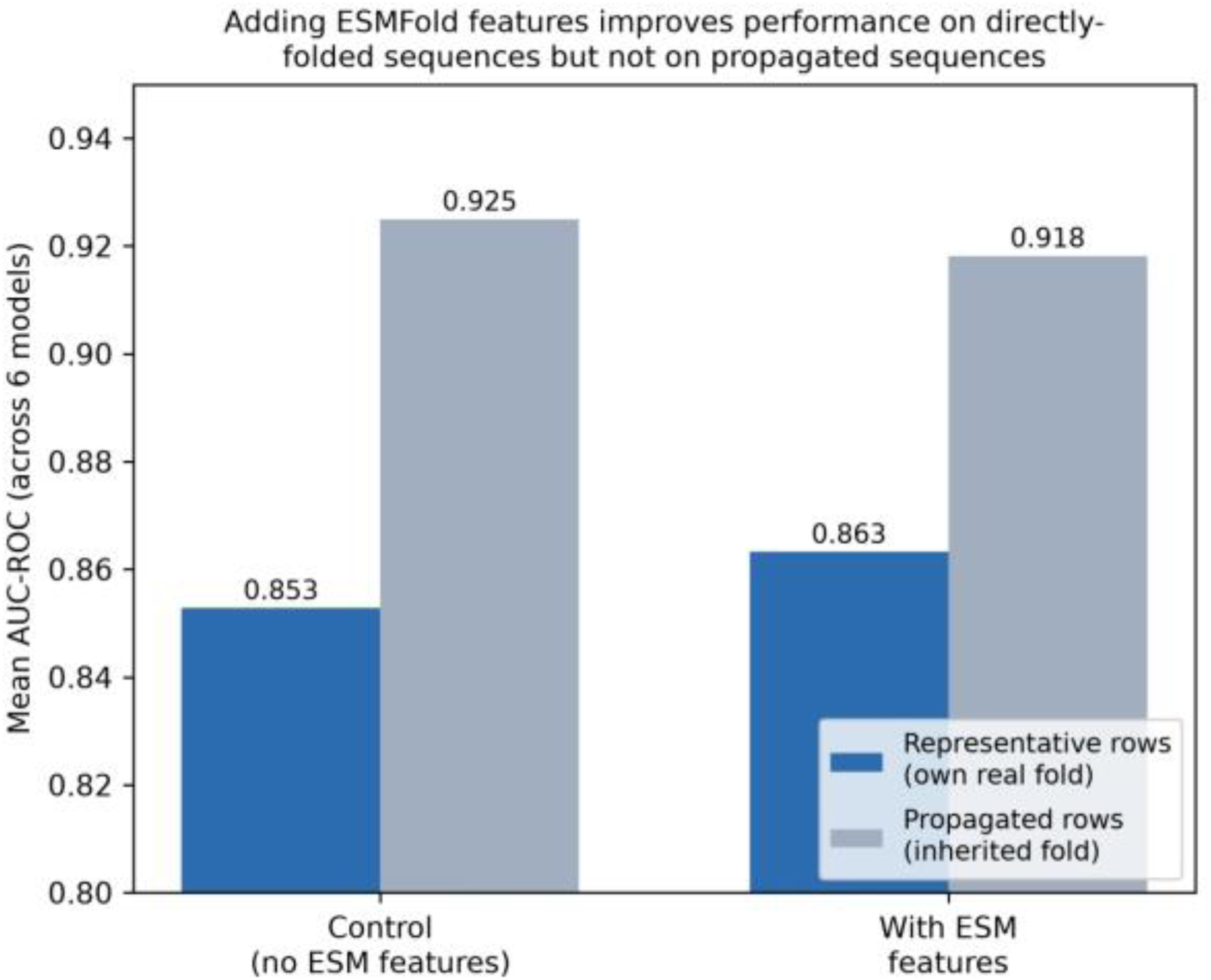
ESMFold propagation diagnostic. Mean AUC-ROC (across 6 models) for test sequences with their own computed fold (representative) versus inherited fold (propagated), with and without ESMFold features.

Comparing models with and without ESMFold features within each subset removes differences between the subsets, including length, from the comparison. Within the subsets, the differences were in the direction originally predicted by the propagation-noise hypothesis. On representative sequences, where ESMFold features reflect a real, individually computed fold, adding these descriptors raised the AUC from 0.853 to 0.863, a difference of +0.010. Conversely, on propagated sequences, where ESMFold features are inherited rather than computed, their inclusion lowered the AUC from 0.925 to 0.918, a drop of -0.007. Because propagated sequences outnumber representatives approximately six to one in the test set, this small negative difference is likely to dominate the aggregate ablation metrics reported in Table 3. Consequently, the aggregate ablation metrics alone may obscure these subset-level differences, which could give the impression that structural features are uniformly uninformative.

Figure 3 shows this decomposition using four bars grouped into two pairs. The first pair compares representative-sequence AUC-ROC values without and with ESMFold features (0.853 versus 0.863), a positive shift that may reflect a small signal in individually computed structures. The second pair compares propagated-sequence AUC-ROC values without and with ESMFold features (0.925 versus 0.918), a negative shift that may reflect a cost of inheriting a neighbor’s structural values rather than computing unique ones. Read together, the four bars separate two groups of sequences that a single aggregate AUC-ROC value conflates. These differences point in opposite directions and are consistent with the small net aggregate performance loss reported in Table 3.

These results are compatible with individually computed structures adding a small amount of predictive value, but the +0.010 AUC difference was observed on only 376 sequences, was not tested for significance, and may lie within sampling variability. An alternative explanation for the small decrease on propagated sequences is that identical propagated feature values within each training cluster act as cluster identifiers that the models can overfit, rather than as noisy structural estimates; we did not distinguish between these explanations. In addition, ESMFold predictions and pLDDT values are less reliable for very short peptides, which make up much of this dataset, and the predicted structures describe isolated monomers rather than membrane-bound conformations. The propagation approximation was adopted specifically to make folding computationally tractable under the constrained 2-core CPU environment without GPU access, rather than reflecting an evaluation that structure is unhelpful. This approach may dilute any structural benefit in the aggregate, which matters for future extensions of this work.

The limitation is a specific, addressable consequence of the computational budget available for this study, rather than evidence against incorporating structural features in antimicrobial peptide machine learning pipelines more broadly. With sufficient computational resources to fold every individual sequence rather than propagating features from cluster representatives, the aggregate ablation result might shift toward the +0.010-signal demonstrated on representative sequences, rather than the small net loss produced by the propagation approximation reported in Table 3.

### Interpretability

SHAP analysis of the optimal architecture, specifically the Random Forest classifier trained on sequence composition and physicochemical features, identified net charge features (pcp_Charge and pcp_ChargeDensity) as the highest-ranked predictors (Table 4, Fig. 4). This finding matches the early charge verification step reported during dataset construction. The agreement between a pre-modeling group-mean comparison and a post-hoc tree-ensemble attribution is consistent with a charge-driven classification signal, but both analyses use the same labeled dataset, so the agreement shows that charge is the dominant difference between these positives and negatives rather than independently confirming mechanism. The negative class was drawn from reviewed proteins with secretory and antimicrobial-annotated entries excluded, which may accentuate this difference.

**Fig. 4.**
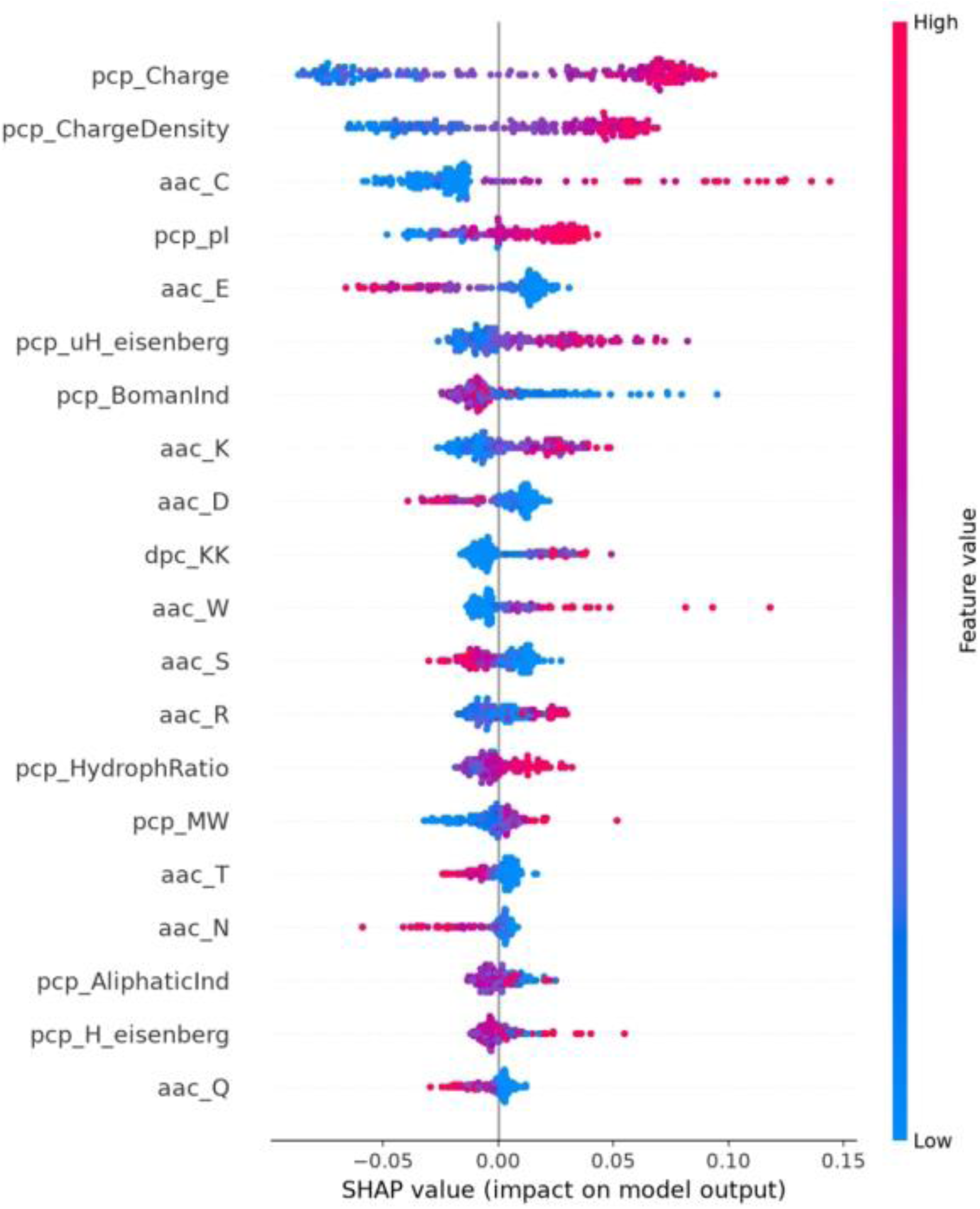
SHAP summary plot for the best model (Random Forest, sequence + physicochemical features), top 20 features

Cysteine composition (aac_C) ranked third, with a positive association: higher cysteine content pushed predictions toward the antimicrobial class (Spearman ρ = 0.47). The effect was uneven. Most sequences contain no cysteine (the median fraction was zero in both classes), but the mean cysteine fraction was higher among antimicrobial sequences than among negatives (0.048 versus 0.020), and 39.6% of antimicrobial sequences versus 32.1% of negatives contained at least one cysteine, so the signal is carried by a minority of cysteine-rich sequences. This is consistent with disulfide-stabilized peptide families being present among the positive sequences, but we did not test this explanation. Cysteine content is not part of the cationic-amphipathic mechanism captured by the other leading features.

The isoelectric point (pcp_pI) ranked fourth with a positive association (ρ = 0.92), and glutamate composition (aac_E) ranked fifth with a negative association (ρ = −0.85), so higher glutamate content pushed predictions away from the antimicrobial class. Both are consistent with a charge-driven mechanism, in which acidic residues lower net charge. Because a peptide’s isoelectric point is a direct function of its charged-residue composition, its presence in the ranking reflects an expected correlation with the charge features rather than an independent signal. Hydrophobic moment (pcp_uH_eisenberg) ranked sixth.

This descriptor detects periodicity in residue hydrophobicity along a sequence [29], which serves as the structural signature of an amphipathic helix where hydrophobic and hydrophilic residues alternate in a pattern that concentrates each type on opposite faces of the folded structure. Its appearance in the top ten, alongside charge metrics, recovers the two defining properties of the classical membrane-disrupting AMP mechanism, namely net cationic charge and amphipathic helical geometry, without either constraint being hard-coded into the model rules.

The Boman index (pcp_BomanInd), ranked seventh, is a composite descriptor proposed as a proxy for a peptide’s potential for protein–protein interactions [28]. It had a negative association (ρ = −0.59): higher values pushed predictions away from the antimicrobial class. Because it aggregates per-residue values, it likely overlaps with the charge and hydrophobicity features, and we do not interpret it as an independent mechanism.

Cationic-residue features closed out the top ten through lysine composition (aac_K, rank 8) and the paired-lysine dipeptide (dpc_KK, rank 10), both exhibiting a positive SHAP direction. Lysine is one of the two standard residues that carries a full positive charge at physiological pH. Its appearance both individually and as a repeated dipeptide motif serves as a secondary, residue-level confirmation of the charge mechanism visible in the aggregate charge descriptors at the apex of the ranking.

Aspartate composition (aac_D) ranked ninth with a negative association (ρ = −0.80), mirroring glutamate: higher aspartate content pushed predictions away from the antimicrobial class, as expected for an acidic residue that lowers net charge.

Taken together, eight of the top ten features relate directly to charge or amphipathicity: net charge, charge density, isoelectric point, glutamate, aspartate, hydrophobic moment, lysine and the lysine–lysine dipeptide. In every case the direction matched the cationic-amphipathic model: features that raise net positive charge pushed predictions toward the antimicrobial class, and acidic residues pushed them away. The remaining two, cysteine content and the Boman index, are not obviously part of that mechanism. Mean |SHAP| values at ranks 7 and 8 (0.0150 and 0.0149) and at ranks 9 and 10 (0.0120 and 0.0118) are nearly identical and were estimated on only 200 sequences, so the composition of the lower half of the top ten should be treated as approximate.

### Gram-positive/Gram-negative Spectrum

Beyond overall antimicrobial classification, the pipeline was evaluated to determine if it generalizes a related but distinct prediction task: activity spectrum against Gram-positive versus Gram-negative bacteria. The two bacterial classes differ substantially in envelope architecture. Gram-positive bacteria present a thick, relatively porous peptidoglycan layer, while Gram-negative bacteria are bounded by an additional outer membrane rich in lipopolysaccharide. This structural variance has long been recognized as a primary factor shaping which AMPs are active against which class [6, 7].

Out of 2,876 cleaned, spectrum-annotated peptides, 88.2% were active against Gram-positive bacteria, 83.3% against Gram-negative, and 71.5% against both. Because most spectrum-active peptides demonstrate efficacy against both classes rather than exclusively one, this was treated as a multi-label task consisting of two independent binary targets, rather than forcing a single mutually exclusive category per peptide. Collapsing the two into a single three-class target (Gram-positive only, Gram-negative only or both) would have fragmented an already imbalanced label distribution into smaller, noisier classes without changing the underlying biology being modeled. The reported percentages imply that every peptide in this subset was active against at least one group, and a label of “not active” against a group may reflect absence of testing or annotation rather than demonstrated inactivity.

Random Forest again performed best for both targets, trained on the same sequence-composition and physicochemical feature space used for the primary classification task, with no dedicated structural or envelope-specific features engineered for this task specifically. The 2,876 spectrum-annotated peptides were split into 2,289 training / 587 test, following the same cluster-disjoint procedure used for the primary task. Gram-positive activity was predicted at AUC-ROC 0.8712 (MCC 0.5890); Gram-negative activity at AUC-ROC 0.8507 (MCC 0.4817) (Table 5).

**Table 5.** Gram-spectrum Secondary Task - Best Model per Target.

| **Target** | **Model** | **AUC-ROC** | **AUPRC** | **F1** | **MCC** |
|---|---|---|---|---|---|
| Gram-positive active | Random Forest | 0.8712 | 0.9801 | 0.9630 | 0.5890 |
| Gram-negative active | Random Forest | 0.8507 | 0.9655 | 0.9254 | 0.4817 |

The AUC-ROC gap between the two targets is modest (0.021), but the MCC gap is larger (0.107) and worth separating from any biological interpretation before reaching for one. Two explanations are plausible, neither of which is empirically confirmed. First, MCC is more sensitive to class-prevalence differences than AUC-ROC, and the annotated dataset carries a real prevalence difference between targets, with 88.2% Gram-positive-active versus 83.3% Gram-negative-active. This difference could by itself compress the Gram-negative MCC relative to the Gram-positive MCC at a similar underlying discrimination ability.

Verification of this effect was attempted by recovering the per-class confusion-matrix counts from the saved evaluation outputs, but because the pipeline’s results tables record only aggregate metrics (AUC-ROC, AUPRC, F1-score, MCC) rather than raw prediction counts, this explanation remains an untested hypothesis. Second, a biological contribution is also plausible in principle. The Gram-negative outer membrane is a structurally distinct barrier from the Gram-positive peptidoglycan layer. If it selects for AMP properties beyond what sequence composition and standard physicochemical descriptors capture, a feature space with no structural tier could underperform specifically on the Gram-negative target. Because this pipeline’s feature set does not include a per-target structural comparison, both explanations are reported as open hypotheses rather than demonstrated causes, and a full accounting of this gap is deferred to future work that logs per-class prediction counts.

### Effect of Split Design and a Charge-Only Baseline

Across 20 repeated splits (seeds 42 to 61), the Random Forest reached a mean AUC-ROC of 0.945 ± 0.004 under random splitting and 0.923 ± 0.012 under cluster-disjoint splitting (range 0.898–0.941). Random splitting gave a higher AUC-ROC in all 20 repetitions, with a mean paired difference of 0.022 (SD 0.012; range 0.004–0.048; 95% CI of the mean 0.016–0.027; Table 6). The primary cluster-disjoint split (seed 42, AUC-ROC 0.939) ranked second of the 20 and therefore lies near the upper end of the distribution; the mean across splits (0.923) is the more conservative summary. A logistic regression on net charge, charge density and isoelectric point reached a mean AUC-ROC of 0.793 ± 0.017 under cluster-disjoint splitting (0.817 on the primary split), and raw net charge used directly as a score gave 0.813 on the primary split. The full-feature Random Forest therefore added about 0.13 AUC-ROC over the charge features (mean difference 0.131).

**Table 6.**
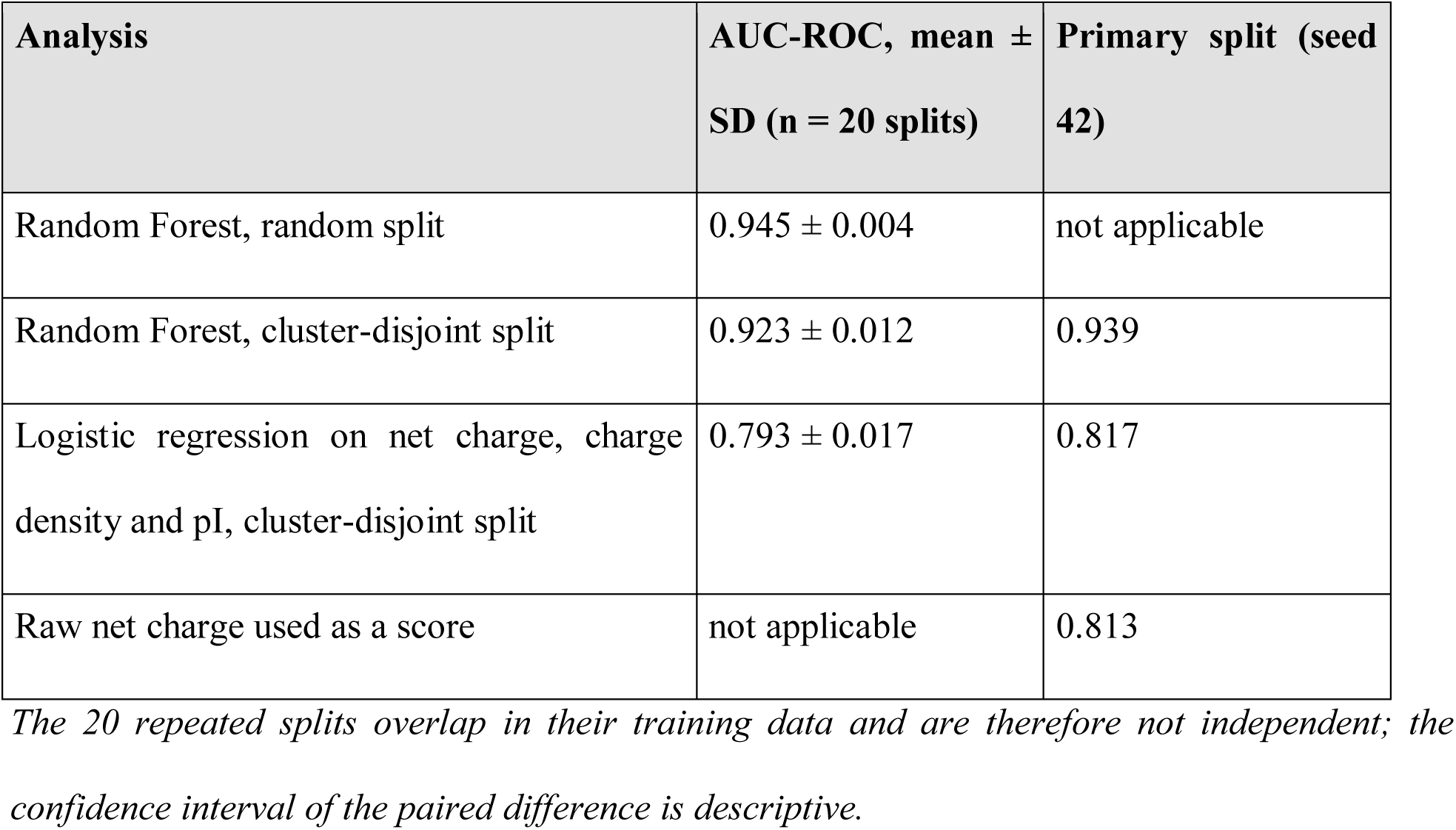
Split Design and Charge Baseline.

### External Check: Scoring of Clinical-Stage Peptides

The clinical-development dataset from the DRAMP, containing 96 peptides that reached preclinical or clinical trials [10], was scored with the trained model as a qualitative check. Forty entries were excluded for non-standard residues and one for being shorter than five residues, applying the same cleaning rules as for the training data, which left 38 peptides. Sixteen of these 38 (eight in the training set and eight in the test set) had an exact sequence match in the general dataset under a different DRAMP identifier, including LL-37, protegrin-1 and nisin A, and were removed. The remaining 22 candidates have no exact-sequence overlap with the data used to build or evaluate the model; this screen was based on sequence identity and did not exclude close homologs (see Discussion). For these 22 peptides the mean predicted probability was 0.73 and the median was 0.82; 18 scored at least 0.5, 12 at least 0.7 and 8 at least 0.9 (Fig. 5). Because the set contains only clinical-stage peptides and no negatives, no AUC can be computed, and Random Forest probabilities are not calibrated, so these values should be read descriptively.

**Fig. 5.**
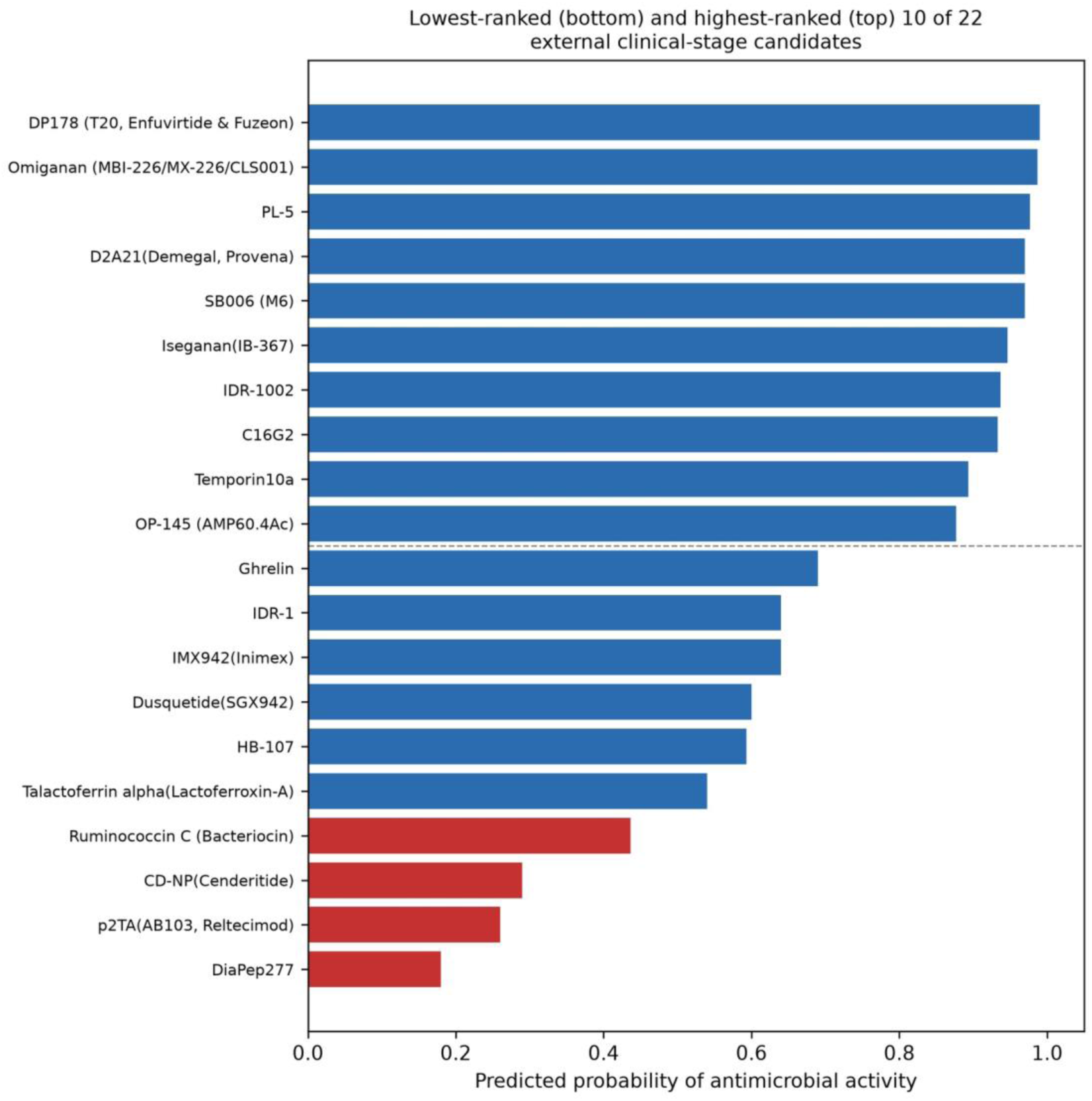
Predicted antimicrobial-activity probability for the top and bottom 10 of 22 clinical-stage candidate peptides with no exact-sequence match in the training or test sets.

Enfuvirtide scored highest at 0.99, followed by omiganan (0.99) and PL-5 (0.98). Omiganan, iseganan and D2A21, which also rank in the top six, are cationic membrane-active peptides. This distribution is consistent with the model recognizing the charge-and-amphipathicity signature learned from the training data (Table 4, Fig. 4) in peptides whose mechanisms match that signature. Because the screen removed only identical sequences, scores for peptides that are close homologs of training sequences may be inflated by similarity to the training data.

One case within the high-scoring group is highly instructive. Enfuvirtide, an HIV fusion inhibitor with no antibacterial indication, scored 0.99, the highest of the 22 candidates. This outcome is not interpreted as a straightforward model error. Enfuvirtide is a 36-residue peptide derived from the HR2 region of HIV-1 gp41. It blocks viral fusion by binding the HR1 region of gp41, and it contains a tryptophan-rich segment that interacts with lipid membranes. It therefore shares helical, membrane-interacting features with membrane-active AMPs without sharing their bactericidal mechanism. This case illustrates a limitation of the feature space and the model constructed on it. The feature set captures generalized amphipathic-helix and membrane-interaction signatures rather than an antimicrobial mechanism specifically. Consequently, a peptide optimized for viral-fusion interference can trigger the same learned signature as a peptide optimized for bacterial membrane lysis. This constraint bears directly on how any score from this model should be interpreted for a truly novel candidate rather than an already-characterized peptide.

The lowest-scoring candidates were informative and sharpen the contrast with the Enfuvirtide case. DiaPep277 (0.18) is an immunomodulatory peptide derived from heat-shock protein 60, developed for type 1 diabetes, which acts on immune cell signaling rather than a membrane target. Reltecimod (0.26) is a CD28 receptor-mimetic peptide that modulates T-cell co-stimulatory signaling to dampen the hyperinflammatory response in severe infection, developed for necrotizing soft-tissue infections and sepsis-associated organ dysfunction, representing an immunomodulatory mechanism rather than an antibacterial one. Cenderitide (0.29) is a chimeric natriuretic peptide that acts through particulate guanylyl cyclase receptors on cardiac and vascular tissue, reflecting a receptor-mediated signaling mechanism with no membrane-disruptive component.

All three are listed in DRAMP’s clinical-development dataset, yet none acts through membrane disruption, and the model scored all three low, as would be expected if it has learned a membrane-interaction signature. Read alongside the enfuvirtide case, this contrast is consistent with the interpretation that the model has learned a representation closer to “cationic amphipathic membrane-interacting peptide” than “antimicrobial peptide” in the full clinical sense, and it illustrates the distinction between biophysical mechanism and categorical labels.

## Discussion

### Principal Findings and Biophysical Interpretations

This pipeline’s central result is not simply that a Random Forest trained on sequence composition and physicochemical descriptors reaches AUC-ROC 0.939 on a cluster-disjoint test set; it is that this number was obtained under an evaluation designed to prevent two failure modes documented in the AMP-ML literature: negative-class sampling bias [19] and, less commonly addressed, positive-class redundancy leakage. Cluster-level, group-stratified splitting after CD-HIT clustering at 40% identity reduces a specific shortcut a model could otherwise exploit: recognizing a near-duplicate SAR-series sibling rather than generalizing from sequence properties. The resulting AUC-ROC (0.939, 95% CI 0.931–0.948) and MCC (0.671, 95% CI 0.643–0.707) fall within the range reported for other AMP classifiers (Table 7), although the evaluation designs differ and the figures are not directly comparable. The confidence intervals resample test sequences rather than clusters and are conditional on this partition, so the spread across repeated partitions (SD 0.012) is a more realistic indication of uncertainty. These headline figures come from one cluster-disjoint partition (seed 42) that ranked second of 20 repeated partitions, and the model and feature tier were selected on the same test set. Across 20 repeated cluster-disjoint splits the mean AUC-ROC was 0.923 ± 0.012, and random splitting raised AUC-ROC in every repetition (mean 0.022, range 0.004–0.048; Table 6), so the inflation attributable to sequence redundancy was consistent but modest on this metric. Threshold-dependent metrics such as MCC may be more sensitive, which we did not test. A logistic regression on charge features alone reached a mean AUC-ROC of 0.793, so the full feature set added about 0.13 AUC-ROC over charge.

**Table 7.** Evaluation-Paradigm Comparison with Prior AMP-ML Studies.

| **Study** | **Dataset (size & source)** | **Evaluation split paradigm** | **Homology control** | **Reported headline metric(s)** |
|---|---|---|---|---|
| This study | 13,469 peptides (DRAMP v5.0 + UniProt) | Cluster-disjoint (CD-HIT-clustered, group-stratified) | CD-HIT, 40% identity, both classes independently | AUC-ROC 0.939, MCC 0.671 |
| Bhadra et al. [13] | 3,268 AMP + 166,791 non-AMP | 10-fold cross-validation, 1:3 pos:neg ratio | Not reported | AUC-ROC ~0.99, MCC ~0.9 |
| Meher et al. [12] | 3,417 antibacterial positives (Xiao benchmark) | 10-fold cross-validation | Not reported | Accuracy 93.84%, MCC 0.88 |
| Yan et al. [14] | 188 sequences, <=30 aa, balanced | Not stated in source | Not reported | AUC-ROC ~0.85, accuracy ~77% |
| Bhangu et al. [18] | 394-peptide held-out test set | Held-out test set; construction not detailed | Not reported | Accuracy ~87% (XGBoost classifier) |
| Ghulam et al. [16] | Study-specific benchmark | Not stated in source | Not reported | Accuracy 97.29% (dipeptide-composition features; 84.42% with amino-acid composition); AUC-ROC 98.91% |

*Based on the methodologies reported in the referenced studies, none described cluster-disjoint train/test splitting. Some of these benchmarks may nevertheless have been redundancy-reduced by sequence-identity filtering before splitting, a related but different control that we did not assess for each study. Where no such control was applied, the reported metrics may be optimistic relative to a cluster-disjoint evaluation of the same models, for the same reason that a leakage-prone split inflates apparent performance. Table 7 does not represent a performance leaderboard; rather, these metrics were produced under fundamentally different evaluation constraints and cannot be compared on a common footing. The AUC-ROC of 0.939 in this study was obtained under a strict cluster-disjoint protocol, which the studies in Table 7 did not report using. It is shown for context and is not evidence of outperforming them.*

The SHAP analysis is consistent with a mechanism the field already expects. Net cationic charge was the dominant driver, and eight of the top ten features relate to charge or amphipathicity, with acidic residues (glutamate, aspartate) pushing predictions away from the antimicrobial class (Table 4, Fig. 4). This recovers the classical cationic-amphipathic-helix model of membrane disruption through a method that had no access to that mechanism as prior knowledge. That a simple pre-modeling group-mean comparison and a post-hoc attribution on the trained model both identify charge as the dominant signal suggests the finding is a property of the data and not of how the SHAP explainer was configured. Both analyses use the same labels, however, and the negative class (reviewed proteins with secretory and antimicrobial-annotated entries excluded) may accentuate the charge difference; net charge alone, used as a score, reached an AUC-ROC of 0.813 on the primary split. SHAP attributions describe the trained model, not causal biology. The leading feature not explained by charge or amphipathicity is cysteine content, which was positively associated with the antimicrobial class; we suggest this may reflect disulfide-rich peptide families among the positive sequences, but we did not test this.

### The Structural Feature Pipeline Trade-off

A methodologically important result of this study is not a headline number: the ESMFold structural-feature ablation looked unexpected on first inspection, and a closer look changed how it should be interpreted. Table 3’s aggregate ablation shows the ESMFold-inclusive feature tier underperforming sequence-and-physicochemical features for most classifiers. Reported at face value, that result would support a simple but potentially misleading conclusion, namely that real three-dimensional structure adds no predictive value over sequence-derived descriptors for AMP activity prediction. Relying solely on the aggregate numbers in Table 3 would have led to precisely that argument.

The diagnostic decomposition in the ESMFold case-study section shows why that conclusion would have been premature.

CD-HIT’s default behavior of designating the longest sequence in each cluster as its representative created a potential confound: sequences with their own computed fold are systematically longer (mean 47.5 residues) than sequences inheriting a propagated fold (mean 22.4 residues), and sequence length may have its own, structure-independent relationship with classification difficulty in this dataset. A no-ESMFold control run reproduced a similar representative-versus-propagated gap (−0.072 AUC) with no structural features present at all, against −0.055 AUC with them included, indicating that the raw gap is not caused by the structural features. Length is a plausible driver, but we did not test it directly. Comparing models with and without ESMFold features within each subset showed a small positive difference (+0.010 AUC, not tested for significance) on sequences with a computed fold, against a small negative difference (−0.007) on sequences that inherited one from a cluster neighbor.

This distinction is critical. It separates two claims that look identical in an aggregate ablation table but carry opposite implications for future iterations of this work. The assertion that structure does not aid prediction is a biological claim that would argue against investing further effort in structural features for AMP prediction. Conversely, the assertion that structure provides utility but that the propagation approximation adopted to make folding computationally tractable under a constrained 2-core CPU environment without GPU access dilutes the aggregate benefit is an engineering claim. This engineering constraint motivates, rather than argues against, further evaluation of structural features given adequate computational resources to fold every individual sequence rather than a representative subset. Distinguishing between these two claims requires, at minimum, splitting the test set by fold provenance and running the control experiment, and even then our data, which include no significance test for the subset difference, do not settle the question.

The aggregate metric alone can obscure this distinction. A counterintuitive result is a prompt to investigate rather than a conclusion to report, although the investigation here is itself limited by the small subset and the absence of a significance test.

### Methodological Limitations and Functional Confounders

The clearest limitation this paper demonstrates directly, rather than states abstractly, is that the model’s discriminative signal is not specific to the antimicrobial mechanism. The external scoring results make this concrete: enfuvirtide, an HIV fusion inhibitor with no antibacterial function, scored 0.99, the highest score among the 22 external candidates, and in practice indistinguishable from the scores of classical membrane-disrupting peptides such as omiganan (0.99). Enfuvirtide’s helical, membrane-interacting character shares biophysical features with membrane-disrupting AMPs, and the model, trained on sequence composition and physicochemical descriptors dominated by charge and hydrophobic moment (Table 4), appears to have learned a broader category than intended: amphipathic, membrane-interacting peptide, rather than antimicrobial peptide specifically. This is a data-evidenced boundary on what this model and plausibly any classifier trained on a similar feature space without an explicit non-membrane-active-peptide negative class can be trusted to discriminate. It should be read as a direct constraint on interpreting any single score from this model for a genuinely novel candidate: a high score indicates membrane-interaction potential consistent with AMP activity, not a confirmed antimicrobial mechanism.

A second limitation concerns negative-class construction. Sidorczuk et al. [19] demonstrated that AMP-ML benchmark performance is highly sensitive to how the negative class is built, with different sampling strategies producing substantially different apparent accuracy for identical architectures. This pipeline addresses that concern only partially. The random-fragment-extraction strategy used here, paired with an explicit annotation-keyword exclusion filter, is a deliberate, auditable control against the most likely contamination route, reviewed UniProt proteins with real but undocumented antimicrobial activity, but it remains a single negative-sampling paradigm among the eleven catalogued by Sidorczuk et al. [19]. This was an intentional scope boundary for this study rather than oversight. Establishing a genuinely leakage-controlled positive-class evaluation was the primary methodological contribution intended for this research, and testing that contribution’s robustness across multiple negative-sampling regimes represents a substantial additional undertaking. Whether the feature-tier ablation conclusions reported here, particularly the sequence-versus-physicochemical-versus-structural performance plateau in Table 3, hold under alternative negative constructions, such as scrambled-AMP negatives or negatives enriched for high-frequency cytoplasmic sequences, is accordingly a distinct and important direction for future work rather than a question this pipeline answers. In addition, excluding secretory proteins from the negative pool means that negatives differ from AMPs in typical subcellular context, which may exaggerate the apparent importance of charge.

A third limitation concerns the physicochemical feature computation itself. The modlAMP descriptors used in the physicochemical tier [30] assume a uniformly free C-terminus. This assumption does not hold for all natural AMPs: C-terminal amidation is common among membrane-active peptides, and because amidation status is inconsistently or incompletely annotated across DRAMP and comparable repositories, it could not be corrected for systematically in this pipeline.

The resulting error is not random noise distributed evenly across the dataset; it is a systematic charge- and pI-calculation error concentrated specifically in the subset of naturally amidated peptides, a biologically coherent and non-trivial subgroup of the positive class. Given that charge and charge-density features are the two most important predictors identified by SHAP analysis (Table 4), this is a limitation with a plausible, if unmeasured, effect on model performance specifically for amidated sequences, rather than a generic caveat applicable equally to the whole dataset.

A fourth limitation concerns the evaluation design. The headline figures come from one cluster-disjoint partition, the 20 repeated splits share training data and are therefore not independent replicates, and the best model and feature tier were selected on the same test set that is used to report performance, so the headline figures may be slightly optimistic. The bootstrap confidence intervals resample test sequences rather than clusters. The clinical-stage peptide set was screened against the training and test sets only for identical sequences (16 of 38 were removed), so close homologs of training peptides may remain among the 22 candidates, and because every peptide in it is labeled as a clinical-stage AMP, it supports no AUC. Cluster-disjoint splitting at a single 40% CD-HIT threshold also does not rule out leakage from more distant homologs, and we did not measure the maximum sequence identity between test and training sequences. The structural comparison rests on 376 directly folded test sequences and was not tested for significance; structures were predicted through a remote service (the ESM Metagenomic Atlas API) whose underlying model version may change, and ESMFold is less reliable for the very short peptides that make up much of this dataset.

### Comparison with Prior Benchmarks

A core structural constraint must be acknowledged prior to evaluating these results against the broader literature: a direct, head-to-head comparison against other published antimicrobial peptide (AMP) classifiers utilizing identical training and testing partitions was not executed. Consequently, the literature comparisons detailed below must be interpreted with this evaluation boundary firmly in mind. Reported baseline metrics from prior studies were produced under evaluation designs that differ from the current study’s protocol in ways that materially affect the mathematical meaning of a reported number. Most importantly, these differences concern whether the train/test splitting strategy is controlled for sequence redundancy. Table 7 summarizes what each study’s own methodology documents, rather than attempting to rank performance across incompatible evaluation regimes.

Two additional structural differences warrant explicit notation rather than inclusion in the tabular summary. First, AmPEP was developed from a pool of 3,268 AMPs and 166,791 non-AMPs but was evaluated at a 1:3 positive-to-negative ratio (about 25% positive) [13]. Our dataset is about 75% positive, and MCC depends on class prevalence, so the MCC values reported for AmPEP (∼0.9) are not comparable with ours.

Second, Deep-AmPEP30, one of the models in Table 7, exclusively targets short peptides [14]. This bounded length regime features a distinct biophysical profile compared to the full length-distribution dataset utilized in this pipeline. These variables do not invalidate qualitative methodological comparisons; they simply demonstrate that Table 7 serves as a map of evaluation methodologies rather than a competitive scoreboard.

## Conclusions

This paper’s central argument is methodological rather than architectural: a trustworthy AMP-ML benchmark depends less on classifier sophistication than on whether the evaluation is constructed to prevent leakage. Cluster-disjoint splitting at 40% CD-HIT identity, applied independently to both classes, reduces a leakage pathway, positive-class sequence redundancy from structure-activity-relationship libraries, that receives less attention in this literature than negative-sampling bias. The Random Forest built under this constraint reaches AUC-ROC 0.939 (95% CI 0.931–0.948) and MCC 0.671 (95% CI 0.643–0.707) on a single cluster-disjoint split, and a mean AUC-ROC of 0.923 across 20 repeated cluster-disjoint splits; the same model scored about 0.02 AUC-ROC higher under random splitting.

Three findings matter beyond the numbers. First, an aggregate ablation suggested that structure does not help, but a subset analysis showed a possible small positive difference (+0.010 AUC, not tested for significance) on sequences with their own computed fold and a small negative difference on sequences with propagated folds; a length confound introduced by CD-HIT’s representative selection complicates the raw comparison. Second, SHAP analysis was consistent with the cationic-amphipathic mechanism and agreed with a simple pre-modeling charge comparison, although both analyses use the same labeled data. Third, external scoring exposed a limitation: the model discriminates membrane-interacting amphipathic peptides more than antimicrobial mechanisms specifically, as the high score for enfuvirtide illustrates.

The broader implication extends past this pipeline. As AMP-ML moves toward generative candidate design, the classifiers scoring generated candidates inherit whatever evaluation weaknesses trained them. A discriminator trained on a redundancy-contaminated split does not become trustworthy because it is repurposed as a filter for novel sequences; if anything, the stakes of an inflated benchmark rise when its output guides which candidates reach a bench. Peptide informatics built on databases curated as biological repositories, not as machine-learning evaluation sets, cannot assume that unpartitioned train/test splits measure what they claim to measure.

Homology-aware, cluster-disjoint evaluation should not be treated as an optional refinement; for a field increasingly using these models to prioritize experimental candidates, it is a precondition for trusting the screening metrics that guide that prioritization.

The complete pipeline structured as a 29-rule Snakemake workflow with logged data provenance at every step is available as open-source software for anyone who wants to check, reuse, or extend it under that standard.

## Declarations

## Acknowledgements

The authors used Claude (Anthropic) during manuscript preparation to assist with drafting and revising prose across all sections, formatting literature citations, and identifying grammatical inconsistencies or minor numerical typos introduced during manual editing. All scientific content, data, analytical decisions, and conclusions originate solely from the authors’ own pipeline. The authors cross-checked and verified all referenced literature, reviewed all AI-assisted text taking full responsibility for the accuracy and integrity of the entire manuscript.

## Ethics statement

Not applicable. This study involved no human participants, human data, or animal subjects. All data used were publicly available peptide sequence and protein databases (DRAMP, UniProtKB/Swiss-Prot).

## Availability of data and materials

The complete 29-rule Snakemake workflow, source code, documentation, environment configuration (environment.yml), metadata-filtration templates, data-provenance manifest, and all result tables and diagnostic figures are available on the repository: https://github.com/isreal-devs/amp-activity-ml. Raw data sources: DRAMP v5.0 (CC BY 4.0; https://dramp.cpu-bioinfor.org), UniProtKB/Swiss-Prot (https://www.uniprot.org). Structure prediction used Meta AI’s ESM Metagenomic Atlas API (https://esmatlas.com).

## Author contributions

All conception, methodology, software, formal analysis, investigation, data curation, writing, and visualization were performed by I.A.O. J.A.Y. reviewed and validated the codebase and the manuscript and formatted it for submission.

